# Elongation capacity of polyunsaturated fatty acids in the annelid *Platynereis dumerilii*

**DOI:** 10.1101/2024.03.15.584978

**Authors:** Marc Ramos-Llorens, Khalida Bainour, Leonie Adelmann, Francisco Hontoria, Juan C. Navarro, Florian Raible, Óscar Monroig

## Abstract

In animals, elongation of very long-chain fatty acid (Elovl) proteins play pivotal functions in the biosynthesis of fatty acids, including the physiologically essential long-chain polyunsaturated fatty acids (LC-PUFA). Polychaetes have important roles in marine ecosystems, contributing not only to nutrient recycling but also exhibiting a distinctive capacity for biosynthesising LC-PUFA, as emphasised in previous research. In order to expand our current understanding of the LC-PUFA biosynthesis in polychaetes, the present study conducted a thorough molecular and functional characterisation of Elovl occurring in the model organism *Platynereis dumerilii*. In this study, we identify six Elovl proteins in the genome of *P. dumerilii*. The sequence and phylogenetic analyses established that four Elovl, identified as Elovl2/5, Elovl4 (two genes) and Elovl1/7, have putative functions in the LC-PUFA biosynthesis. Functional characterisation in yeast confirmed the roles of these elongases in the LC-PUFA biosynthetic pathways, demonstrating that *P. dumerilii* possesses a varied and functionally diverse complement of Elovl enzymes that, along the enzymatic specificities of previously characterised desaturases, enable *P. dumerilii* to perform all the reactions required for the biosynthesis of the LC-PUFA. Importantly, we uncovered that one of the two Elovl4-encoding genes from *P. dumerilii* is remarkably long in comparison with any other animals’ Elovl, which contains a C terminal KH domain unique among Elovl enzymes. The distinctive expression pattern of this protein in photoreceptors strongly suggests a central role in vision.

## 1. Introduction

Long-chain (C_20-24_) polyunsaturated fatty acids (LC-PUFA) play critical roles in various biological functions within animals (Sargent et al. 1995; Arts et al. 2001). Among the most physiologically important LC-PUFA are the omega-3 (n-3) eicosapentaenoic acid (EPA) and docosahexaenoic acid (DHA), and the omega-6 (n-6) arachidonic acid (ARA). These compounds are integral components of cell membranes, influencing membrane fluidity and permeability. LC-PUFA are precursors of signalling molecules termed eicosanoids, which participate in immune responses and inflammation regulation (Dyall et al. 2022). Generally, ARA-derived eicosanoids have pro-inflammatory properties, whereas those from EPA and DHA are anti-inflammatory, playing roles in immune regulation and contributing to cardiovascular health (Lee et al. 2009; Swanson et al. 2012; Manson et al. 2019; Schulze et al. 2020; Harwood, 2023). Additionally, DHA, particularly abundant in brain and retina cell membranes, contributes to neural development and function, supporting cognitive processes and visual acuity (Jeffrey et al. 2001; Janssen et al. 2014; Lauritzen et al. 2016). Fish and seafood are important sources of the n-3 LC-PUFA (EPA and DHA) in the human diet since the production of these essential nutrients occurs primarily in marine ecosystems. Importantly, LC-PUFA are not just beneficial for humans but also play pivotal roles in ensuring the well-being and growth of marine organisms themselves, highlighting the ecological and nutritional significance of LC-PUFA in marine ecosystems (Arts et al. 2001; Twining et al. 2020).

Microorganisms such as marine microalgae, bacteria and other cellular organisms positioned at the bottom of marine trophic webs have been historically regarded as the unique primary producers of n-3 LC-PUFA in the ocean. Upper trophic level organisms such as fish have some capacity to bioconvert short-chain (C_18_) polyunsaturated fatty acids (PUFA) to the C_20-22_ LC-PUFA (EPA, DHA and ARA) (Monroig et al. 2018), but mostly accumulate LC-PUFA by consuming LC-PUFA-rich preys including invertebrates (Arts et al. 2001; Gladyshev et al. 2013; Twining et al. 2020). While invertebrates can acquire LC-PUFA from the diet, recently collected evidence has established that certain groups can biosynthesise C_18_ PUFA de novo, an ability believed to be absent in animals (Kabeya et al. 2018, 2020, 2021; Boyen et al. 2023). Such a metabolic capacity is accounted for by the presence of methyl-end desaturases in Cnidaria, Rotifera, Mollusca, Arthropoda, Nematoda and Annelida (Kabeya et al. 2018). Thus, a methyl-end desaturase with Δ12 desaturase activity enables the biosynthesis of most structurally simple PUFA, i.e., linoleic acid (18:2n-6, LA), which can be subsequently desaturated by a methyl-end desaturase with Δ15 activity to α-linolenic acid (18:3n-3, ALA). The C_18_ PUFA LA and ALA serve as precursors for the biosynthesis of n-6 and n-3 LC-PUFA, respectively. This process involves the activity of specific enzymes, namely front-end desaturases and elongation of very long-chain fatty acids (Elovl) proteins. Like methyl-end desaturases, front-end desaturases introduce double bonds into a fatty acid that acts as a precursor (Sperling et al. 2003). Moreover, Elovl catalyse the condensation step of the fatty acid elongation pathway, resulting in the extension of the fatty acyl chain of a precursor in two carbons (Leonard et al. 2004; Jakobsson et al. 2006; Gregory et al. 2011). Collectively, animals with complementary enzymatic activities contained within methyl-end desaturases, front-end desaturases and Elovl have the potential to biosynthesise the biologically important LC-PUFA and, hence, contribute to the primary production of these essential nutrients.

Polychaetes (Annelida) have been previously demonstrated to possess methyl-end desaturases (Liu et al. 2017; Kabeya et al. 2018, 2020) and front-end desaturases (Ramos-Llorens et al. 2023a). However, to the best of our knowledge, no Elovl have been previously characterised in polychaetes, so it remains uncertain whether they have complete pathways for LC-PUFA biosynthesis. Elovl enzymes vary in their substrate specificity and, among the eight Elovl identified in vertebrates (Elovl1-8) (Monroig et al. 2022), Elovl1, Elovl3, Elovl6 and Elovl7 are generally regarded as elongases with affinity towards saturated and monounsaturated fatty acids, whereas Elovl2, Elovl4, Elovl5 and Elovl8 elongate PUFA substrates and therefore play relevant roles in the LC-PUFA biosynthesis (Leonard et al. 2004; Jakobsson et al. 2006; Castro et al. 2016; Monroig et al. 2022). In invertebrates, several Elovl involved in the LC-PUFA biosynthetic pathways have been characterised in molluscs, crustaceans and echinoderms (Monroig et al. 2012; 2016a; 2016b; 2017; Liu et al. 2013; Kabeya et al. 2017; 2021; Ribes-Navarro et al. 2021; Ramos-Llorens, et al. 2023b; Boyen et al. 2023). Most of the Elovl characterised from invertebrates have been termed as Elovl1/7, Elovl2/5, Elovl4 or Elovl8 according to their phylogenetic relationship with vertebrates’ Elovl, as well as ‘novel’ Elovl when such a relationship could not be established (Monroig and Kabeya, 2018; Monroig et al. 2022). Arguably, the most commonly studied PUFA Elovl from invertebrates are Elovl2/5 and Elovl4 (Monroig et al. 2022). Elovl2/5, an ancestral enzyme that evolved in two distinct genes (Elovl2 and Elovl5) in vertebrates (Monroig et al. 2016b), has been only confirmed to occur in molluscs (Monroig et al. 2016a; Monroig et al. 2017; Ran et al. 2019) and amphioxus (Monroig et al. 2016b). Moreover, Elovl4 are PUFA elongases with well-established roles in the biosynthesis of LC-PUFA, as well as the so-called very long-chain (>C_24_) polyunsaturated fatty acids (VLC-PUFA) (Monroig et al. 2010), key components of cell membranes in photoreceptors of vertebrates (Monroig et al. 2010, Deák et al. 2019). The potential involvement of invertebrates’ Elovl4 in photoreception remains unexplored, with existing studies predominantly focussed on elucidating the functions of the encoded enzymes (Monroig et al. 2018).

The polychaete annelid *Platynereis dumerilii* has emerged as a model organism in the fields of evolution, chronobiology, and developmental biology. Multiple genetic and molecular tools, including a draft genome, have been developed for *P. dumerilii* in recent years (Zantke et al. 2014), providing a unique opportunity for unravelling fundamental biological processes (Özpolat et al. 2021). *P. dumerilii* is an excellent choice as a reference model for LC-PUFA biosynthesis and its regulation because it possesses two functional methyl-end desaturases (Kabeya et al. 2018), as well as two further front-end desaturases (Ramos-Llorens et al. 2023a). While the activities reported for these enzymes cover all the requirements for desaturase-mediated reactions involved in the LC-PUFA biosynthetic pathways, no data on the complement and function of *P. dumerilii* Elovl enzymes have been yet reported. The present study aimed to uncover the repertoire of Elovl proteins in *P. dumerilii*, providing a comprehensive understanding of the roles and diversity of these enzymes in LC-PUFA biosynthesis within the broader context of annelid polychaetes. Moreover, *P. dumerilii* has a variety of photoreceptors, both in the head and along the body axis (Arendt et al. 2004; Backfisch et al. 2013; Randel et al. 2013; Zantke et al. 2013; Ayers et al. 2018; Poehn et al. 2022). Since vertebrate photoreceptors contain VLC-PUFA (Agbaga et al. 2010) and Elovl4 is a key enzyme in the VLC-PUFA biosynthesis (Deák et al. 2019), we analysed the spatial expression of the *P. dumerilii elovl4* to clarify putative roles of the invertebrate Elovl4 in photoreception and vision.

## 2. Materials and methods

### 2.1 Sample collection, RNA extraction and cDNA synthesis

One single *P. dumerilii* individual from an established culture of the inbreed strain “PIN” (Zantke et al. 2014) available at Max Perutz Labs Vienna (Vienna, Austria) was homogenised in TRIzol ™ reagent for further extraction of total RNA according to the manufacturer’s instructions (Thermo Fisher Scientific, Waltham, MA, USA). Complementary DNA (cDNA) was synthesised from 2 µg of total RNA using the Moloney Murine Leukemia Virus Reverse Transcriptase (M-MLV RT; Promega, Madison, WI, USA) using random primers and oligo-dT primers (3:1 mol) (Promega). The obtained cDNA was stored at −20 °C until further use.

### 2.2 Sequence and phylogenetic analysis

Using invertebrate Elovl-like sequences from *Amphimedon queenslandica*, *Artemia franciscana*, *Echinogammarus marinus*, *Octopus vulgaris* and *Tigriopus californicus* as queries (Monroig et al. 2012; Gold et al. 2017; Kabeya et al. 2021; Ribes-Navarro et al. 2021; Ramos-Llorens et al. 2023b), *tblastn* searches were carried out against annelid transcriptomes available at the National Center of Biotechnology Information (NCBI; www.ncbi.nlm.nih.gov [accessed on 11 January 2021]). Additionally, candidate sequences were searched using *blastn* against the draft genome available for *P. dumerilii* (Zantke et al. 2014; Özpolat et al. 2021), in order to validate their authenticity, location and genetic structure with Geneious Prime (v: 2022.1.1; Sievers et al. 2011; Kearse et al. 2012). From the gene localisation results, a schematic figure was generated with the Matplotlib library (v. 3.5.1; Python v.3.8.8; Hunter, 2007). The predicted conserved motives of the putative proteins were subsequently analysed using the *Pfam* online tool (https://pfam.xfam.org/ [accessed on 12 January 2021]), and only sequences containing the complete “*ELO*” domain (PF01151) for Elovl (Mistry et al. 2021) were selected for further analysis. In addition, other functional motifs predicted in the sequences, if any, were analysed. The deduced amino acid sequences of the putative *P. dumerilii* Elovl sequences (six in total and termed as “EloA-F”) were aligned with Clustal Omega (default settings) as implemented in the software Geneious Prime. Characteristic features of Elovl enzymes, including conserved motives distinguishing between PUFA and non-PUFA Elovl were analysed according to Hashimoto et al. (2008). In the case of EloC, due to its uniqueness, studies of the topology of the protein in its localisation in the membrane were carried out using DeepTMHMM (v.1.0.24; Hallgren et al. 2022).

A phylogenetic tree comparing the amino acid sequences of the *P. dumerilii* elongases with homologues retrieved from a variety of vertebrate and invertebrate species was carried out using the maximum-likelihood (ML) method (Whelan et al. 2001). Representatives for all Elovl subfamilies (Elovl1-8), as well as “novel” Elovl were selected for analysis (Xie et al. 2021; Monroig et al. 2022). Briefly, the amino acid sequences were first aligned with MAFFT (Katoh et al. 2013). Then, the alignment was processed with trimAl (Capella-Gutiérrez et al. 2009) as a previous step to generate an ML phylogenetic tree using PhyML (Guidon et al. 2010), and an automated model selection implemented in PhyML based on heuristic strategies with the SMS software (Lefort et al. 2017). The confidence in the resulting phylogenetic tree branch topology in which the associated taxa clustered together in the bootstrap test (1000 replicates) was calculated (Lemoine et al. 2018). The resulting tree was processed in Netwick format (Junier et al. 2010) and visualised using MEGA11 (Tamura et al. 2021).

### 2.3 Molecular cloning of full-length P. dumerilii elongase genes

A selection of the *elovl* candidate genes retrieved from *P. dumerilii* were cloned into the yeast expression vector pYES2 (Thermo Fisher Scientific) for further functional characterisation (below). Selection was based on the presence of specific motives for PUFA elongases (Hashimoto et al. 2008), and the phylogenetics results supporting their close relationship with well-established PUFA elongase families (Castro et al. 2016; Monroig et al. 2022). Four distinct Elovl-like sequences (EloA-D) were identified as putative PUFA Elovl in *P. dumerilii*. Full-length open reading frames (ORF) sequences of the selected putative PUFA elongases from *P. dumerilii* were amplified by PCR (Phusion® DNA polymerase, Thermo Fisher Scientific) with primers containing restriction enzyme sites for further cloning into pYES2 (**Error! Reference source not found.**). The PCR amplification parameters for both *eloA* and *eloB*, were as follows: an initial denaturing step at 98 °C for 3 min, 40 cycles of denaturation at 98 °C for 30 s, annealing at 55 °C for 30 s, extension at 72 °C for 45 s, and a final extension at 72 °C for 10 min. For *eloC*, a two-round PCR strategy was required to amplify the ORF using a high-fidelity PrimeStar ® DNA polymerase (Takara Europe, Saint-Germain-en-Laye, France). The first round of the nested PCR was carried out using the above cDNA as the template with primers designed in the 5’- and 3’-untranslated regions (UTR) of the *eloC* sequence (Table S1). The second round (nested) PCR was performed with primers containing restriction sites as described above (Table S1), and using the first round PCR product as template. Both the first round and nested PCR consisted of an initial denaturing at 98 °C for 3 min, followed by 40 cycles of denaturation at 98 °C for 10 s, annealing at 58 °C for 15 s, extension at 68 °C for 2 min, and a final extension at 68 °C for 10 min. PCR products containing the ORF sequences of the targeted genes (*eloA*, *eloB* and *eloC*) were purified on an agarose gel using the Wizard ® SV gel and PCR clean-up system purification kit (Promega). After restriction with the corresponding restriction enzymes (New England Biolabs, Ipswich, MA, USA) (Table S1) and purification as above, the PCR products were ligated (T4 DNA ligase, Promega) into similarly restricted pYES2. Finally, the full-length ORF sequence of *eloD* was ordered as a synthetic gene (Genscript Biotech, Leiden, The Netherlands), and subcloned into pYES2 as described above. Ligations of the restricted ORF fragments into the pYES2 vector were transformed into competent *Escherichia coli* TOP10™ strain (Invitrogen) and grown in Luria-Bertani (LB) agar plates with ampicillin (100 μg/mL). Subsequently, colonies were screened by PCR, and potential positive clones were grown overnight in LB media for plasmid production and further purification with GenElute™ Plasmid Miniprep Kit (Sigma-Aldrich). After confirmation of the insert sequence by sequencing (DNA Sequencing Unit, IBMCP-UPV, Valencia, Spain), the constructs pYES2-PDEloA, pYES2-PDEloB, pYES2-PDEloC and pYES2-PDEloD were obtained.

### 2.4 Functional characterisation by heterologous expression in yeast

Each of the plasmid constructs (pYES2-PDEloA, pYES2-PDEloB, pYES2-PDEloC and pYES2-PDEloD) were transformed into *Saccharomyces cerevisiae* INV*Sc*1 competent cells (Thermo Fisher Scientific) using the *S.c.* EasyComp® Transformation kit (Thermo Fisher Scientific). Yeast selection was carried out on *S. cerevisiae* minimal medium minus uracil (SCMM^−ura^) agar plates for 3 d at 30 °C. Next, one single colony from each construct transformation was incubated in SCMM^-ura^ broth for 2 d at 30 °C and under constant shaking (250 rpm) to generate a yeast culture with an OD_600_ of 8–10. Then, an aliquot of the yeast cultures was inoculated in 5 mL of SCMM^-ura^ broth contained in individual 150 mL Erlenmeyer flasks to provide an OD_600_ of 0.4. Culture from each flask was grown for 4 h at 30 °C and under constant shaking until they reached an OD_600_ of ∼1, at which point the transgene expression was induced by supplementing the culture media with galactose at 2% (w/v). In addition to galactose, an aliquot of the following potential PUFA substrates was exogenously supplemented in individual flasks: α-linolenic acid (18:3n-3), linoleic acid (18:2n-6), stearidonic acid (18:4n-3), γ-linolenic acid (18:3n-6), eicosapentaenoic acid (20:5n-3), arachidonic acid (20:4n-6), docosapentaenoic acid (22:5n-3) and docosatetraenoic acid (22:4n-6). Final concentrations of the exogenously supplied PUFA substrates were 0.5 mM (C_18_), 0.75 mM (C_20_), and 1.0 mM (C_22_) to compensate for decreased efficiency of uptake with increased chain length (Lopes-Marques et al. 2017). All FA substrates (>98–99% pure) used for the functional characterisation assays were obtained from Nu-Chek Prep, Inc. (Elysian, MN, USA). Yeast culture reagents, including galactose, nitrogen base, raffinose, tergitol NP-40, stearidonic acid (>99%) and uracil dropout medium were obtained from Sigma-Aldrich (St. Louis, MO, USA). The yeast cultures were maintained at 30 °C and under constant shaking (250 rpm) for 2 d until yeast was harvested by centrifugation at 1500 g for 2 min. Yeast pellets were washed twice with 5 mL of ddH_2_O, homogenised in 6 mL of 2:1 (v/v) chloroform:methanol containing 0.01% (w/v) butylated hydroxytoluene (BHT, Sigma-Aldrich) as antioxidant, and stored at −20 °C for a minimum of 24 h in an oxygen-free atmosphere until further lipid extraction.

### 2.5 Fatty acid analysis from yeast

The determination of the fatty acid composition of the recombinant yeast expressing the *P. dumerilii elovl* was performed following the methodologies described by Ribes-Navarro et al. (2021). Briefly, total lipids were extracted from the homogenised yeast samples following Folch’s method (Folch et al. 1956). Subsequently, total lipids were used to prepare fatty acid methyl esters (FAME) through acid-catalysed transmethylation (Christie, 1982), and purified using thin-layer chromatography on 20 x 20 cm plates (Merk, Darmstadt, Germany). FAME samples prepared from the yeast assays were analysed by gas chromatography (Thermo Trace GC Ultra, Thermo Electron Corporation, Waltham, MA, USA) using a capillary column (30 m × 0.25 mm × 0.25 µm film thickness, TR-WAX, Teknokroma, Spain) and flame ionisation detection (FID). Helium was used as a carrier gas. When further confirmation of the peak identity was required, samples were injected into analysis by an Agilent 6850 Gas Chromatograph system coupled to a 5975 series MSD (Agilent Technologies, Santa Clara, CA, USA) equipped with a Sapiens-5MS (30 m × 0.25 µm × 0.25 µm) capillary column (Teknokroma). The elongation of exogenously supplemented PUFA substrates by the *P. dumerilii* EloA, EloB, EloC and EloD was calculated by the proportion of substrate fatty acid converted to elongated fatty acid product(s) as [product areas/ (product areas + substrate area)]× 100.

### 2.6 In situ hybridisation chain reaction (HCR)

Preliminary analysis of single-cell sequencing revealed that, among the four functionally characterised elongases, only *eloB* and *eloC* showed a tissue-specific expression pattern (Milivojev et al. unpublished results), and were hence considered for in situ hybridisation chain reaction (HCR) analysis. Wild-type premature “PIN” *P. dumerilii* individuals (∼ 10) from the Max Perutz Labs Marine Animal Facility (Vienna, Austria) (Zantke et al. 2014) were anesthetised in a 1:1 (v/v) mixture of 7.5% (w/v) MgCl_2_ and artificial seawater (Tropic Marin, Germany). Worms were decapitated with a cut at a position corresponding to ∼2/3 segments after the first parapodia. The HCR-based multi-labelling protocol for visualising the spatial expression patterns of the *P. dumerilii* EloB and EloC, and cell nuclei, was established according to Ćorić et al. (2023). Briefly, samples were fixed with 1 mL 4% (v/v) paraformaldehyde with phosphate-buffered saline with Tween20 (PTW) (0.1%, v/v, Tween20/phosphate-buffered saline) (Sigma-Aldrich), and subsequently, dehydrated by a series of washing steps with an increase of methanol concentration diluted in PTW buffer. Samples were stored at −20 °C for further processing. Where needed, samples were rehydrated with a decreasing methanol concentration diluted in PTW buffer, and treated with proteinase K (Sigma-Aldrich) for 5 min. Then, samples were pre-incubated with hybridisation buffer at 37 °C for 1 h before adding the corresponding probe mix with hybridisation buffer. After overnight incubation at 37 °C, samples were washed to remove unattached probes several times with probe wash buffer, sodium chloride-sodium citrate buffer and finally, with PTW buffer. Next, amplification buffer was added, and separately, hairpin mixes were prepared with hairpin pairs, namely, the pair B2H1 and B2H2 (EloB) and the pair B1H1 and B1H2 (EloC). Additionally, Hoechst counterstain (Invitrogen) was included in the final mix for cell nuclei staining. Hairpin mixes were added to the amplification buffer containing the samples and incubated overnight at room temperature, protected from light. On the following day, samples were washed several times with sodium chloride-sodium citrate buffer and prepared for microscopy visualisation using a mounting medium. Hybridisation, probe wash and amplification buffers were prepared according to Ćorić et al. (2023). Probe hybridisation buffer, probe wash buffer, amplification buffer, and fluorescent HCR hairpins were purchased from Molecular Instruments (Los Angeles, CA, USA). Hairpins associated with the B1 initiator sequence were labelled with Alexa Fluor 546, and the hairpins associated with the B2 initiator sequence were labelled with Alexa Fluor 647. Other chemicals used for HCR were purchased from Sigma-Aldrich. From Özpolat lab, insitu_probe_generator script (version 0.3.2; https://github.com/rwnull/insitu_probe_generator) was used to generate DNA oligo 3.0-style probes for HCR (Choi et al. 2018; Kuehn et al. 2022; Ćorić et al. 2023) specific to *P. dumerilii eloB* and *eloC* mRNA visualisation (Table S2).

### 2.7 Confocal microscopy visualisation

The anterior parts (heads) of immature adult *P. dumerilii* were used to investigate the spatial expression patterns of the *elovl4*-like genes (*eloB* and *eloC*) with putative roles in photoreception, since they contain brain and eyes where vertebrates’ *elovl4* have been shown to be highly expressed (Monroig et al. 2010, Yeboah et al. 2021). Visualisation of target gene mRNA expression in worm heads was performed on a Zeiss LSM700 confocal microscope (Carl Zeiss, Jena, Germany), as described before (Ćorić et al. 2023). Image analysis was carried out using Fiji/ImageJ software (Shindelin et al. 2012).

## 3. Results

### 3.1 Phylogeny and sequence analysis of the P. dumerilii elongases

A total of six *elovl*-like transcript sequences (EloA-F) were retrieved through extensive search in *P. dumerilii* available ‘omic databases. The phylogenetic trees of the deduce amino acid sequences of the *P. dumerilii* elongases are shown in Fig. 1. The *P. dumerilii* EloA clustered together Elovl2/5 from other invertebrates including molluscs and amphioxus, as well as Elovl2 and Elovl5 sequences from vertebrates (Fig. 1). Moreover, the amino acid sequences of the *P. dumerilii* EloB and EloC grouped along Elovl4 sequences from other animal species including invertebrates and vertebrates (Fig. 1). Additionally, EloD clustered together with the so-called “novel” elongase group, an Elovl1/7-like group, including sequences from crustaceans such as *A. franciscana*, *E. marinus* and *T. californicus*, which have been characterised as PUFA elongases (Ribes-Navarro et al. 2021; Boyen et al. 2023; Ramos-Llorens et al. 2023b) (Fig. 1). Finally, the *P. dumerilii* EloE and EloF grouped together with vertebrate Elovl3 and Elovl6, elongases with no roles on PUFA elongation (Guillou et al. 2010), and were therefore excluded from functional analysis (Fig. 1).

**Figure 1.**
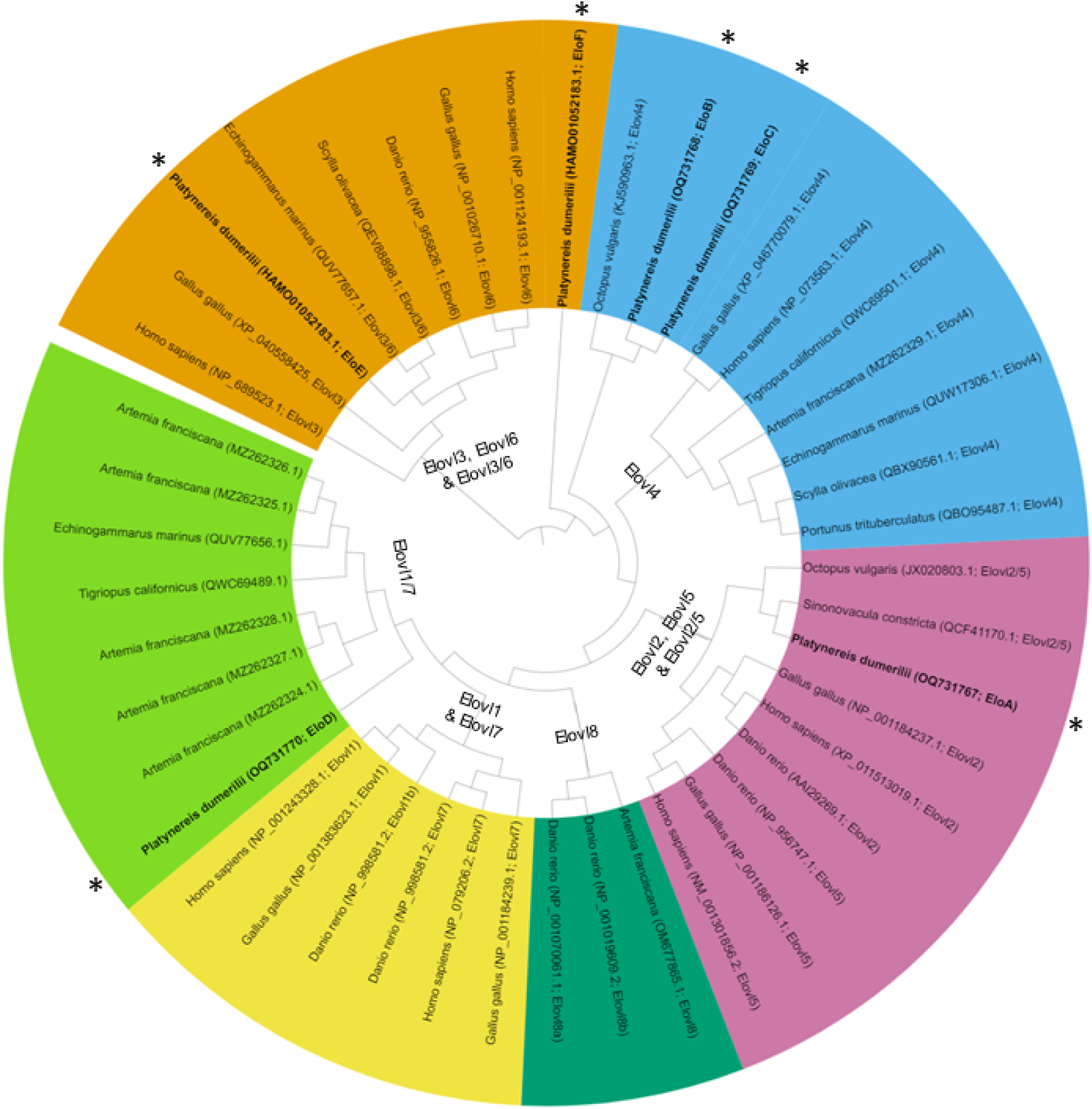
Phylogenetic tree comparing the annelid Elovl enzymes characterised in the present study (asterisks) with other animal elongases. The tree represents the clusters that include Elovl3, Elovl6 and Elovl3/6 enzymes (orange background); Elovl4 enzymes (blue background); Elovl2, Elovl5 and Elovl2/5 enzymes (pink background); Elovl8 enzymes (dark green background), Elovl1 and Elovl7 (yellow background); and Elovl1/7 enzymes (also reported as “novel Elovl”) (light green background).

The amino acid sequences of the *P. dumerilii* EloA-D fulfilled the criteria established by Hashimoto et al. (2008) for PUFA elongases. In detail, the *P. dumerilii* EloA-D contained the distinctive H-box (HXXHH, specifically HV(F/Y)HH) and contained a glutamine (Q) at position −5 from the conserved H-box (Fig. S1). The ORF of the *P. dumerilii* EloA, EloB and EloD contain 927, 936 and 858 base pairs (bp), respectively, encoding putative Elovl proteins of 308, 311 and 285 amino acids. Notably, the ORF of *P. dumerilii eloC* was 2196 bp long, encoding a putative protein of 713 amino acids that, along the ELO motif, includes a KH domain typically found in nucleic acid-binding proteins (Fig. 2A; Fig. S2). In short, the larger length is given by a greater extension of the C terminus where the KH domain is located, which is different from the rest of the elongases analysed. The N terminus is homologous to the rest of the elongases with the “ELO” motif in a similar location (Fig. 2A). This section of the C terminus was predicted to be located inside the cell, which would correspond to the cellular cytosol (Fig. S3). The analysis of the exon-intron structure and the genome location showed that both *eloB* and *eloC* are located in the same genomic neighbourhood (scaffold 47) and have an 8 exon-structure with markedly different lengths of their intronic regions (Fig. 2B). Moreover, the ORF of *P. dumerilii eloA* and *eloD* transcripts contain 6 and 7 exons, respectively (Fig S2), and were located sequentially in the same genomic region (scaffold 1 of the draft genome). The sequences of the *P. dumerilii* elongases studied herein were deposited into the NCBI GenBank with the accession numbers OQ731767 (EloA), OQ731768 (EloB), OQ731769 (EloC) and OQ731770 (EloD).

**Figure 2.**
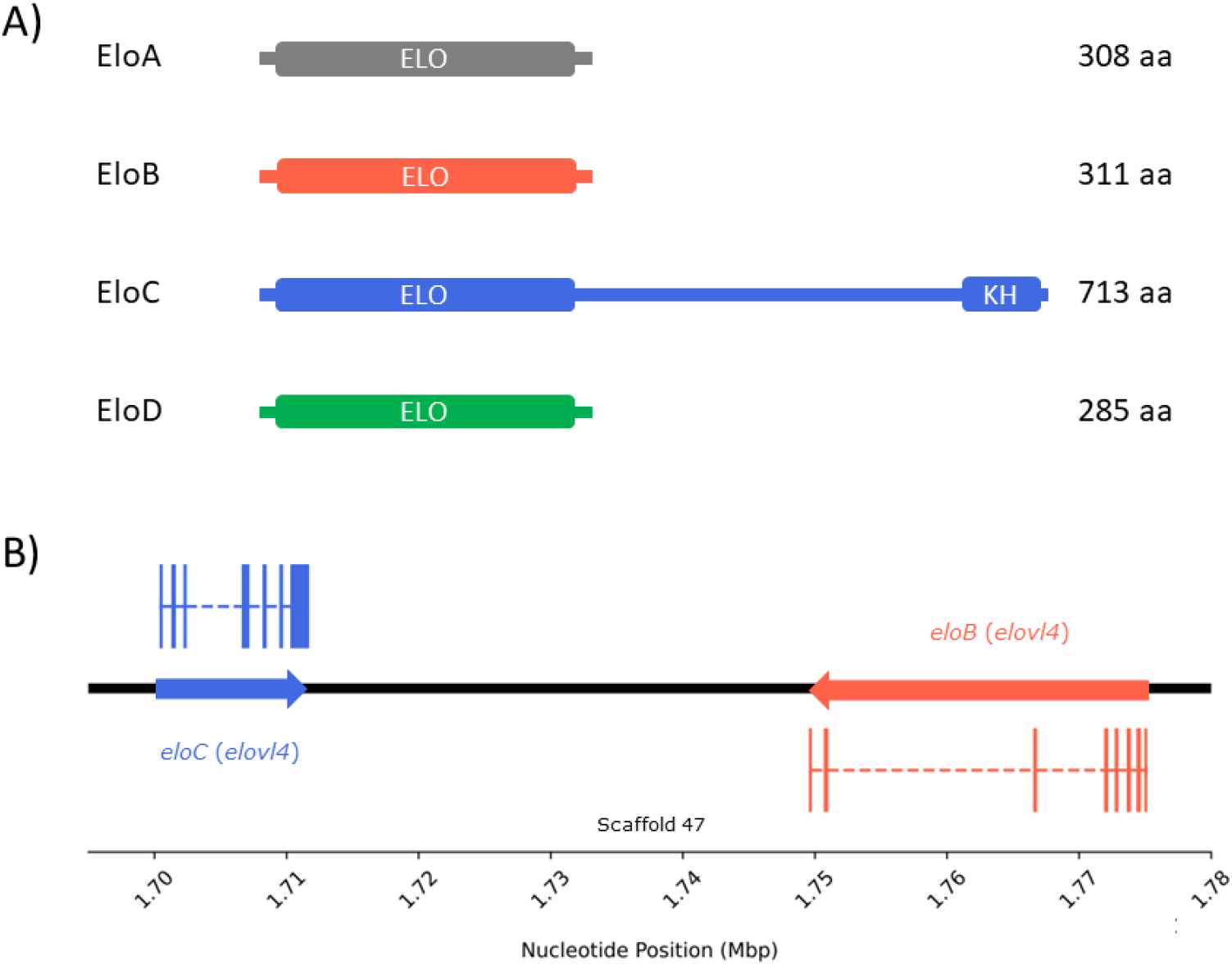
A) Diagram of the predicted functional motifs of the *Platynereis dumerilii* enzymes studied in the present study (EloA-D), as well as their respective sizes in amino acids (aa). The “ELO” motif, associated with the fatty acyl elongase enzyme family (Elovl), is common to all organisms with the presence of these enzymes (Monroig et al. 2022), whereas the KH domain had not been previously described in any Elovl. B) Gene structure comparison of *elovl4* homologues. This representation emphasises the conservation and variation in gene structure between the two homologues within the same scaffold.

### 3.2 Functions of the P. dumerilii elongases in LC-PUFA biosynthesis

The ability of the *P. dumerilii* EloA, EloB, EloC and EloD enzymes for PUFA elongation was assessed by incubating transgenic yeast expressing their ORF in the presence of potential PUFA substrates (18:3n-3, 18:2n-6, 18:4n-3, 18:3n-6, 20:5n-3, 20:4n-6, 22:5n-3 and 22:4n-6). All four *P. dumerilii* Elovl enzymes were functionally active towards a range of PUFA substrates but with varied preferences (Table 1). EloA was able to elongate the four C_18_ PUFA substrates tested (18:3n-3, 18:2n-6, 18:4n-3 and 18:3n-6) to their respective C_20_ elongation products (20:3n-3, 20:2n-6, 20:4n-3 and 20:3n-6), but not over C_20_ and C_22_ PUFA substrates (Table 1). For the *P. dumerilii* EloB, our results showed that this elongase was able to convert all C_18_ PUFA substrates to C_20_ products, whereas the C_20_ PUFA substrates were elongated to the corresponding C_22_ products (Table 1). More specifically, the *P. dumerilii* EloB catalysed the conversion of 18:3n-3, 18:2n-6, 18:4n-3and 18:3n-6, to 20:3n-3, 20:2n-6, 20:4n-3 and 20:3n-6, respectively (Table 1). Moreover, the *P. dumerilii* EloB elongated the C_20_ 20:5n-3 and 20:4n-6 to the C_22_ products 22:5n-3 and 22:4n-6, respectively. Similar results were obtained for the *P. dumerilii* EloD that, like EloB, elongated all the C_18_ and C_20_ PUFA assayed but showed no activity towards C_22_ substrates (Table 1). The functional characterisation of the *P. dumerilii* EloC revealed distinctive traits when compared to EloA, EloB and EloD. Unlike the other PUFA elongases from *P. dumerilii*, EloC showed activity towards all assayed substrates, including not only C_18_ and C_20_ PUFA, but also C_22_ substrates like 22:5n-3 and 22:4n-6 (Table 1). Moreover, while EloA, EloB and EloD produced one single +2C product, conversions by the *P. dumerilii* EloC resulted in step-wise elongation products up to C_24_ (marked with an “*” in Table 1). No apparent activity towards the yeast endogenous saturated and monounsaturated fatty acids was detected for any of the functionally characterised *P. dumerilii* elongases.

**Table 1.** Functional characterisation of the PUFA elongases from *Platynereis dumerilii*. Conversions of exogenously supplemented polyunsaturated fatty acid (FA) substrates were calculated according to the formula [product areas/ (product areas + substrate area)] x 100.

| FA substrate | Product | EloA | EloB | EloC | EloD | Activity |
| --- | --- | --- | --- | --- | --- | --- |
| 18:3n-3 | 20:3n-3 | 0.51 | 3.46 | 19.82* | 2.66 | C<sub>18</sub> to C<sub>20</sub> |
| 18:2n-6 | 20:2n-6 | 0.31 | 0.68 | 5.81* | 1.99 | C<sub>18</sub> to C<sub>20</sub> |
| 18:4n-3 | 20:4n-3 | 0.38 | 2.70 | 13.41* | 6.35 | C<sub>18</sub> to C<sub>20</sub> |
| 18:3n-6 | 20:3n-6 | 0.38 | 1.79 | 9.52* | 2.91 | C<sub>18</sub> to C<sub>20</sub> |
| 20:5n-3 | 22:5n-3 | n.d. | 0.61 | 2.23* | 3.08 | C<sub>20</sub> to C<sub>22</sub> |
| 20:4n-6 | 22:4n-6 | n.d. | 0.58 | 1.95* | 0.74 | C<sub>20</sub> to C<sub>22</sub> |
| 22:5n-3 | 24:5n-3 | n.d. | n.d. | 0.21 | 1.93 | C<sub>22</sub> to C<sub>24</sub> |
| 22:4n-6 | 24:4n-6 | n.d. | n.d. | 0.16 | 0.90 | C<sub>22</sub> to C<sub>24</sub> |

n.d., not detected; \*, Conversions of step-wise reactions are included.

### 3.4 Spatial expression of the P. dumerilii elovl4 elongases

The spatial expression patterns of the *P. dumerilii elovl4*-like sequences (*eloB* and *eloC*) were investigated by in situ HCR. Expression of *eloB* transcripts was found to be scattered in the adult brain (Fig. 3a). The expression of *eloC* was most prominently detected in the *P. dumerilii* adult eyes (Fig. 3b). However, we also observed a co-labelling of *eloC* with *eloB* in brain cells (arrowheads in Fig. 3c). The Supplementary Figures S4-S7 show more details on the expression patterns of the *P. dumerilii eloB* and *eloC* in the anterior part of the body, as well as the control samples.

**Figure 3.**
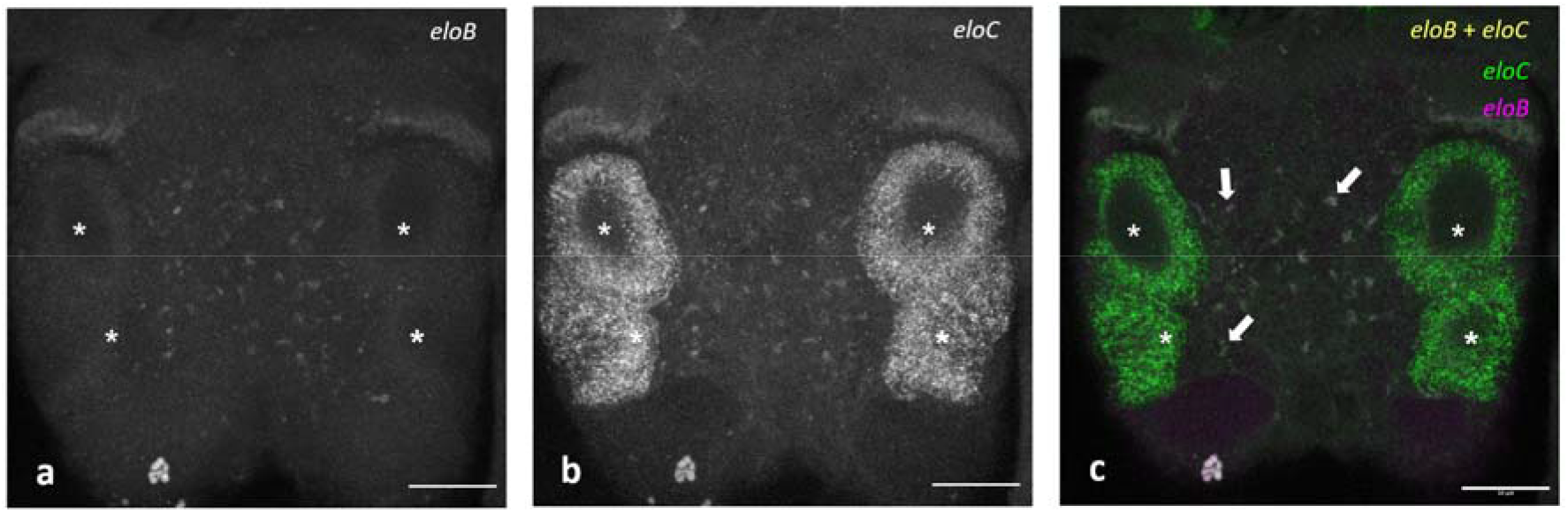
Visualisation, by in situ Hybridisation Chain Reaction (in situ HCR), of the expression of *eloB* and *eloC* in the *Platynereis dumerilii* head: (**a**) Visualisation of *eloB* probes, (**b**) Visualisation of *eloC* probes, and (**c**) Co-visualisation of *eloC* probes (green), *eloB* probes (red) and co-localisation of *eloB* and *eloC* probes (yellow). Arrowheads in panel C show some co-localisation sites of *eloB* and *eloC* probes. Adult eyes are marked with asterisks. Acquisitions by confocal microscopy; scale bar = 50 μm.

## 4. Discussion

The capacity of an animal for LC-PUFA biosynthesis depends upon the gene complement of fatty acyl desaturases and elongases existing in that species, as well as the substrate specificities of the encoded enzymes. This study aimed to improve our comprehension of LC-PUFA biosynthesis in polychaetes by conducting a thorough molecular and functional characterisation of Elovl. Specifically, the study focused on delineating the roles of Elovl in the LC-PUFA biosynthetic pathways within the model organism *P. dumerilii*, complementing previous investigations on methyl-end desaturases (Kabeya et al. 2018) and front-end desaturases (Ramos-Llorens et al. 2023a) (Fig. 4).

**Figure 4.**
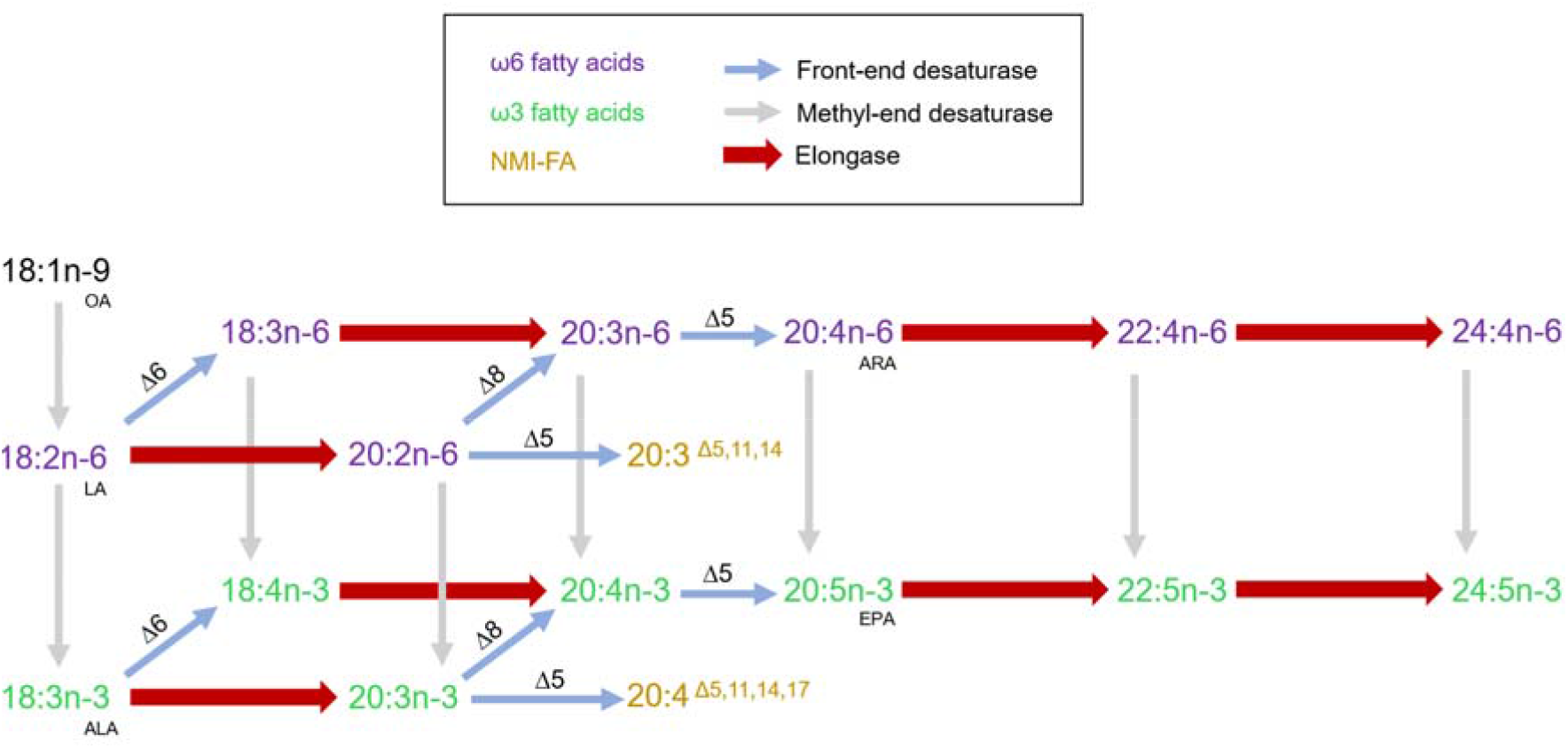
An illustration of the PUFA and LC-PUFA biosynthetic pathways demonstrated in *Platynereis dumerilii*. Reactions catalysed by the characterised *P. dumerilii* enzymes (EloA, EloB, EloC and EloD) are marked with wide red arrows. The desaturase reactions (thin arrows) were characterised previously in *P. dumerilii* by Kabeya et al. (2018) in the case of the methyl-end desaturase enzymes and Ramos-Llorens et al. (2023a) in the case of the front- end desaturase enzymes. OA, oleic acid; LA, linoleic acid; ALA, α-linolenic acid; EPA, eicosapentaenoic acid; ARA, arachidonic acid; NMI-FA, non-methylene interrupted fatty acid.

The Elovl sequence retrieval strategy used in the present study revealed that *P. dumerilii* has six distinct *elovl* genes (EloA-F). The analysis of the deduce amino acid protein sequences of the so-called “EloE” and “EloF” showed that these Elovl do not satisfy with all the characteristics shared among PUFA elongases (Hashimoto et al. 2008; Nie et al. 2021). Consistently, the phylogenetic analysis clearly established that the *P. dumerilii* EloE and EloF clustered with Elovl3 and Elovl6, enzymes with roles in the elongation of non-PUFA substrates (Guillou et al. 2010). These results support the idea that the *P. dumerilii* EloE and EloF do not play relevant functions in LC-PUFA biosynthesis and were, therefore, excluded from further analysis. More interestingly, the sequence and phylogenetic analyses of the other four Elovl identified in *P. dumerilii* (EloA-D) strongly suggested putative roles of these enzymes in the LC-PUFA biosynthetic pathways. Indeed, the *P. dumerilii* EloA can be categorised as an Elovl2/5, an ancestral protein occurring in certain groups of invertebrates that gives origin to two distinct PUFA Elovl in vertebrates, namely Elovl2 and Elovl5 (Monroig et al. 2016b). In addition to the amphioxus *Branchiostoma lanceolatum*, Elovl2/5 has been only reported in molluscs, including the cephalopods *O. vulgaris* (Monroig et al. 2012) and *Sepia officinalis* (Monroig et al. 2016a), and the bivalves *Crassostrea angulata* (Zhang et al. 2018) and *Mimachlamys nobilis* (Liu et al. 2013). It is interesting to note that while the amphioxus Elovl2/5 can elongate C_18_, C_20_ and C_22_ PUFA substrates, and the mollusc Elovl2/5 can elongate C_18_ and C_20_ PUFA, functional assays performed on the *P. dumerilii* EloA confirmed this enzyme has a more restricted range of elongation substrates since it was only capable of elongating C_18_ substrates but not C_20_ or C_22_. Since the herein Elovl2/5 from *P. dumerilii* is the first enzyme of this type that has been functionally characterised in annelids, it is unclear whether the substrate specificities reported here are conserved across this animal group. More clearly though, the apparent lack of elongation capacity of the *P. dumerilii* Elovl2/5 towards C_20_ and C_22_ PUFA can be compensated for those of the other *P. dumerilii* PUFA elongases coexisting in this species.

The *P. dumerilii* EloB and EloC classified as Elovl4 according to our phylogenetic analyses, were both able to elongate C_18_ and C_20_ PUFA but, in addition, EloC could also elongate C_22_ PUFA substrates to the corresponding C_24_ products. In invertebrates, the mollusc Elovl4 was able to produce VLC-PUFA up to C_34_ when expressed in yeast (Monroig et al. 2017; Ran et al. 2019). In crustaceans, however, Elovl4 typically produce PUFA of a maximum acyl chain length of C_24_ like the *P. dumerilii* EloC (Kabeya et al. 2021; Ramos-Llorens et al. 2023b) and, occasionally, C_26_ (Sun et al. 2020; Ribes-Navarro et al. 2021; Boyen et al. 2023). The functional analyses of the two *P. dumerilii* Elovl4 do not enable us to clarify whether polychaetes’ Elovl4 can participate in the biosynthesis of VLC-PUFA as occurs in other invertebrates. Nevertheless, the ability to biosynthesise C_24_ PUFA by the *P. dumerilii* EloC but not EloB, together with other distinctive features in both spatial patterns of expression and structural motives of the encoded enzymes, strongly suggested that the two Elovl4-like elongases from *P. dumerilii* are not functionally redundant.

The HCR analysis showed that expression of EloB is scattered in the head region, containing the brain. On the other hand, EloC has a more specific expression pattern in the eye. These cell types are related to photoreception and vision (Rhode, 1992). The expression patterns observed for the two Elovl4 from *P. dumerilii* largely resemble those described in the model teleost zebrafish (*Danio rerio*), whereby the so-called “Elovl4a” had a scattered expression in the head region (neural tissues) like the *P. dumerilii* EloB, whereas the Elovl4b isoform was specifically expressed in photoreceptors like the *P. dumerilii* EloC. Collectively, the consistency in the spatial expression across animals described above highlights the relevant roles of Elovl4 in photoreception, which is largely conserved across animals. In agreement, the sole human *ELOVL4* has been shown to be highly expressed in the retina’s outer nuclear layer, and rod and cone photoreceptor inner segments (Agbaga et al. 2010). As noted above, *P. dumerilii* is used as a model organism in photoreception research due, among other reasons, to its distinctive combination of larval and adult eyes in *P. dumerilii*. Emphasising the exceptional photoreception capabilities of *P. dumerilii*, this study unveils the existence of two bona fide *elovl4* genes, a unique trait among invertebrates, which usually have one sole Elovl4 (Monroig and Kabeya, 2018; Monroig et al. 2022). Whereas the 3R whole genome duplication (WGD) accounts for the occurrence of the abovementioned paralogous Elovl4 in teleosts (a and b isoforms) (Monroig et al. 2010), the results from the present study do not clarify the origin of two Elovl4 genes in *P. dumerilii*. Both genes are in the same scaffold but in different orientations, which gives clues about the genuineness of both Elovl4. However, it was interesting to observe that, while the *P. dumerilii* EloB and EloC share a high degree of identity along their respective ELO motifs, the EloC but not EloB expands for further 1262 bp in its C terminus, which contains the KH domain. Proteins containing KH domains are typically involved in various cellular processes by binding to RNA or single-strand DNA, thus exerting both post-transcriptional and translational regulation (Valverde et al. 2008). Searches for potential homologues of the unusually long Elovl4 from *P. dumerilii* (EloC) in other annelids, including *Capitella teleta*, *Dimorphilus gyrociliatus*, *Eisenia fetida* and *Streblospio benedicti* among others, have yielded no positive hits. These findings imply that the presence of KH domain-containing Elovl4 is relatively limited across annelids with a yet unknown function in coordination with the elongase motif on the protein. This finding, in the context of the localised expression of this unique gene in the ocular tissue, makes us hypothesise about the existence in *P. dumerilii* of some type of regulation or signalling mediated by nucleic acids in which this special type of Elovl4 intervenes.

Completing the PUFA elongase capacity of *P. dumerilii* is EloD, annotated as Elovl1/7 based on our phylogenetics results. The *P. dumerilii* EloD clustered together with previously characterised elongases annotated as Elovl1/7-like or “novel” elongases from crustaceans such as *T. californicus* (Kabeya et al. 2021), *E. marinus* (Ribes-Navarro et al. 2021), *Platychelipus littoralis* (Boyen et al. 2022) and *A. franciscana* (Ramos-Llorens et al. 2023b). The number of this gene type in crustaceans can range from a single copy like in *E. marinus* (Ribes-Navarro et al. 2021), to three copies in *T. californicus* (Kabeya et al. 2021) and five copies in *P. littoralis* and *A. franciscana* (Boyen et al. 2022; Ramos-Llorens et al. 2023b). Except for the so-called “Elovl1d” from *P. littoralis*, whose action led to the production of VLC-PUFA up to C_28_ (Boyen et al. 2022), the crustacean Elovl1/7 enzymes typically recognise C_18_, C_20_ and C_22_ PUFA as substrates that are elongated to the corresponding up to C_24_ products (Ribes-Navarro et al. 2021; Ramos-Llorens et al. 2023b). In agreement, the substrate specificities of the *P. dumerilii* Elovl1/7 (EloD) indicated an affinity of this elongase towards C_18_/C_20_/C_22_ PUFA substrates, which are elongated up to C_24_ polyenes like reported in the majority of crustacean Elovl1/7 that have been functionally characterised so far.

In conclusion, the present study demonstrates that *P. dumerilii* has six Elovl-encoding genes. Four of the *P. dumerilii* Elovl, identified as Elovl2/5, Elovl4 (2 genes) and Elovl1/7, include all the sequence characteristics of PUFA elongases. The functional characterisation assays in yeast confirmed the roles of these elongases in the LC-PUFA biosynthetic pathways. Therefore, our findings unequivocally establish that *P. dumerilii* harbours a varied and functionally diverse set of Elovl enzymes that, together with the activities attributed to previously characterised desaturases (Kabeya et al. 2018; Ramos-Llorens et al. 2023a), enable this annelid to perform all the reactions required for the biosynthesis of the LC-PUFA EPA and ARA. In addition, this study reported for the first time the occurrence of an Elovl4 with a KH domain fused to the Elo motif. The distinctive expression pattern of this protein strongly indicates its likely pivotal role in vision, specifically in photoreception.

## Acknowledgements

We thank Myriam Lizanda Piqueras, Júlia Pérez Ara, Juan G. Haro Blasco, Nadja Milivojev, Lukas Orel and Dunja Rokvic for their invaluable technical support and expertise throughout this research project. Additionally, we thank Andrej Belokurov, Margaryta Borysova, and Netsaneh Getachew for their excellent support at the University of Vienna marine facility. This study was funded through the project IMPROMEGA Agencia Española de Investigación, Spain, grant no. RTI2018-095119-B-100, MCIN/AEI/ FEDER/UE / MCIN/AEI/10.13039/501100011033/ and FEDER “A way to make Europe”. L.A. was supported by the Austrian Science Fund (FWF) DOC 72 grant (SCORPION) (to F.R.) and a PhD fellowship by the German Academic Scholarship Foundation. Additionally, this study forms part of the ThinkInAzul programme and was supported by MCIN with funding from the European Union NextGenerationEU (PRTR-C17.I1) and by Generalitat Valenciana (THINKINAZUL/2021/26). The European Molecular Biology Organization funded MRL with an EMBO Scientific Exchange Grant (no. 9163).

## Data availability

Additional information needed will be shared upon request.

## Author contributions

***Marc Ramos-Llorens***: Investigation, Methodology, Formal analysis, Writing – original draft, Visualisation. ***Khalida Bainour***: Investigation, Methodology, Formal analysis. ***Leonie Adelmann***: Investigation, Methodology, Formal analysis. ***Francisco Hontoria***: Conceptualization, Supervision, Writing – review & editing, Funding acquisition. ***Juan C. Navarro***: Conceptualization, Investigation, Supervision, Writing – review & editing, Funding acquisition. ***Florian Raible***: Conceptualization, Supervision, Writing – review & editing, Funding acquisition. ***Óscar Monroig***: Conceptualization, Supervision, Writing – review & editing, Funding acquisition.

