## Supplementary Materials for "Elongation capacity of polyunsaturated fatty acids in the annelid *Platynereis dumerilii*"

**Supplementary Tables**

**Table S1**. List of primers and their nucleotide sequences used for molecular cloning of
Platynereis dumerilii eloA, eloB and eloC. Restriction sites in primers used for cloning into the yeast expression vector pYES2 are underlined.

| Primer name | Sequence 5'-3' |
| --- | --- |
| PDEloA-For-HindIII | CCCAAGCTTAAGATGGCGCAACTTGC |
| PDEloA-Rev-XbaI | GCTCTAGAACATTAACTGGACTTTTCATGGTGA |
| PD3-For-HindIII | CCCAAGCTTTGTTATCATGGAGGTC |
| PDEloB-Rev-XhoI | CGCTCGAGCGTCAAGTTCACTCTCAC |
| PDEloC-5UTR | AGGCACATGGTGATTTGAAAAGGAGT |
| PDEloC-3UTR | CAATATTAGATCGGGTCGGCCTCT |
| PDEloC-For-BamHI | CCCGGATCCATTATGGATGTCCTGAAAGACACAG |
| PDEloC-Rev-XhoI | CCGCTCGAGTCTGCGCTTTAAATTGTCACTC |

**Table S2**. List of DNA oligo probes for in situ hybridisation chain reaction (in situ HCR) detection of a Platynereis dumerilii eloB and eloC.

| Gene | Probe sequences |
| --- | --- |
| *eloB* | CCTCGTAAATCCTCATCAaaTGCCATTGGCCAGTGGTCGTCCATT  ACTACTTCCTCTTTTTGTTATCTCCaaATCATCCAGTAAACCGCC  CCTCGTAAATCCTCATCAaaTCCATTTTTGCTGATGTAGCCATTT  ATCTCCACTGCCGTTGGCCAACGCTaaATCATCCAGTAAACCGCC  CCTCGTAAATCCTCATCAaaTTCTTTTTGTGCTTCGGCGGGATGT  ATATGGCCGTTTGTGTATTCTGTTTaaATCATCCAGTAAACCGCC  CCTCGTAAATCCTCATCAaaCCAGGATTGAACCTCCGTAGAAAAT  CGTGGAAATAAAAGTTGAGGAAAAGaaATCATCCAGTAAACCGCC  CCTCGTAAATCCTCATCAaaACAGCCGACAATGAGGGACTGGGCA  AGCCCACTGCATCCAGAGAGGGAAGaaATCATCCAGTAAACCGCC  CCTCGTAAATCCTCATCAaaAATTGCATCCTGGTGAGATATCTCT  TGGATCATGCCCGCTACAAACTGGAaaATCATCCAGTAAACCGCC  CCTCGTAAATCCTCATCAaaCCATTGACGCCAGGCCGTAGTATGA  ACCAGAGGTACTTCTGCACTTTGGGaaATCATCCAGTAAACCGCC  CCTCGTAAATCCTCATCAaaCAGGGCTCCAAAAAATGCTTGGCCA  CATCACCACGTGGATCCAAGAGTTCaaATCATCCAGTAAACCGCC  CCTCGTAAATCCTCATCAaaATGGGGAACATGGTCGCGTGATGGT  GGGACCCACTTGACTCCGACCCACCaaATCATCCAGTAAACCGCC  CCTCGTAAATCCTCATCAaaTCTTTCGGAGAATGAAGAAAAACGT  CGTGAAGGAAGGAAATCTGCGTGTTaaATCATCCAGTAAACCGCC  CCTCGTAAATCCTCATCAaaAAACCACCACAACGCTTTTGCTATC  CATGAATTCTAGACATTTGGAAAAGaaATCATCCAGTAAACCGCC  CCTCGTAAATCCTCATCAaaACAAGTTGACAACTATAACTGTAAC  ACTTCATATGGGTTGTCACTGCGATaaATCATCCAGTAAACCGCC  CCTCGTAAATCCTCATCAaaAAAATATATGAAAGTTGAGCAAAAC  GTATTGTAGTACATACTATAAGCTCaaATCATCCAGTAAACCGCC  CCTCGTAAATCCTCATCAaaTCTTAGTTTCATAGGTTCTCGATGC  TAAAGCATTGTATATTACAATGGGAaaATCATCCAGTAAACCGCC  CCTCGTAAATCCTCATCAaaAGAAGATACAAGACTACTCCTGCCC  ATTAGTTTTGGACCTATCCAAACTAaaATCATCCAGTAAACCGCC  CCTCGTAAATCCTCATCAaaAATCTTCTACTCTTTTATCTGCATA  TCTTCCAATAGGCGTCCATCAAAAAaaATCATCCAGTAAACCGCC |
| *eloC* | GAGGAGGGCAGCAAACGGaaACATTCCAAACGACATGCGAACCTT  CCCCATCTTCCTTCCTCTCCTCGCTtaGAAGAGTCTTCCTTTACG  GAGGAGGGCAGCAAACGGaaGATACTTCAAAGGCCTCACTTTCAT  AAAGACTGCTTCAACGGAACAATTGtaGAAGAGTCTTCCTTTACG  GAGGAGGGCAGCAAACGGaaTGCCTTTCAGTTCGTTAGGAATCGA  ATGCGGTAGCACTTGCCGCACTTGCtaGAAGAGTCTTCCTTTACG  GAGGAGGGCAGCAAACGGaaACGGATACAACGGTGGGTCATTTCA  AGTGGCAGCAATTGCCTCTTTGTAGtaGAAGAGTCTTCCTTTACG  GAGGAGGGCAGCAAACGGaaGTAAACTTCAGTTTAATCATAGTAG  CCAGTTACCCAGGCAGATCTGCTTCtaGAAGAGTCTTCCTTTACG  GAGGAGGGCAGCAAACGGaaGATTAAGGAGATCTACTTGAAGTCT  TAGCTATTTGACTGCCCAACATTTTtaGAAGAGTCTTCCTTTACG  GAGGAGGGCAGCAAACGGaaGTAAATTTCTGAAAACTTGCAAGCT  CTTGTTGTTTGGTGGTAGAGTTGTAtaGAAGAGTCTTCCTTTACG  GAGGAGGGCAGCAAACGGaaACAGGAAAGGCCAGTGCATCTTGAA  TCTACGATCATTTTCAGGTCTGAAAtaGAAGAGTCTTCCTTTACG  GAGGAGGGCAGCAAACGGaaAATCGAATGGCTGACACATATCCAT  CTTCTGCGCCAATCATCAATAAGACtaGAAGAGTCTTCCTTTACG  GAGGAGGGCAGCAAACGGaaATCCCATCAAATAGTACTAGCCCTT  GGCAGGCAATCCAAGAGAGCTACAGtaGAAGAGTCTTCCTTTACG  GAGGAGGGCAGCAAACGGaaAGATGATCTAGTCAATTCTAGGTCA  TGCATAATCGCATTCTTTTTCAAGCtaGAAGAGTCTTCCTTTACG  GAGGAGGGCAGCAAACGGaaTCCTGCAACAAGCTAGCGAGCTTCT  TCCATCTCTTCCTTTTCTTTGTTATtaGAAGAGTCTTCCTTTACG  GAGGAGGGCAGCAAACGGaaGTTTGAGACGATCAATATCTCCGTA  TTTCCTTGACTAACGTTTCCATTTCtaGAAGAGTCTTCCTTTACG  GAGGAGGGCAGCAAACGGaaGAGTTGTTGATTTGTGTCATCTTGA  GTTGTCCAATTCTTGACGTATCAGAtaGAAGAGTCTTCCTTTACG  GAGGAGGGCAGCAAACGGaaCTACCACGTTTTTGCTTTTGAGCCT  TCGTCCTTTTTACTTTTGGGTCTACtaGAAGAGTCTTCCTTTACG  GAGGAGGGCAGCAAACGGaaTATCAAATGAGCACATATCTGTTGC  CCGATGTTGCATCGAATGTGGAGTCtaGAAGAGTCTTCCTTTACG  GAGGAGGGCAGCAAACGGaaAGCGGGAGTTTCATCAGTCTCGTCA  TGGATTCCTGGTGCGTTTTGGTCTAtaGAAGAGTCTTCCTTTACG  GAGGAGGGCAGCAAACGGaaGTAACCTGATCTTGGTCTTTCAGTG  ATTTGGTCATAATCTTCTGCGTTTAtaGAAGAGTCTTCCTTTACG |

**Supplementary Figures**

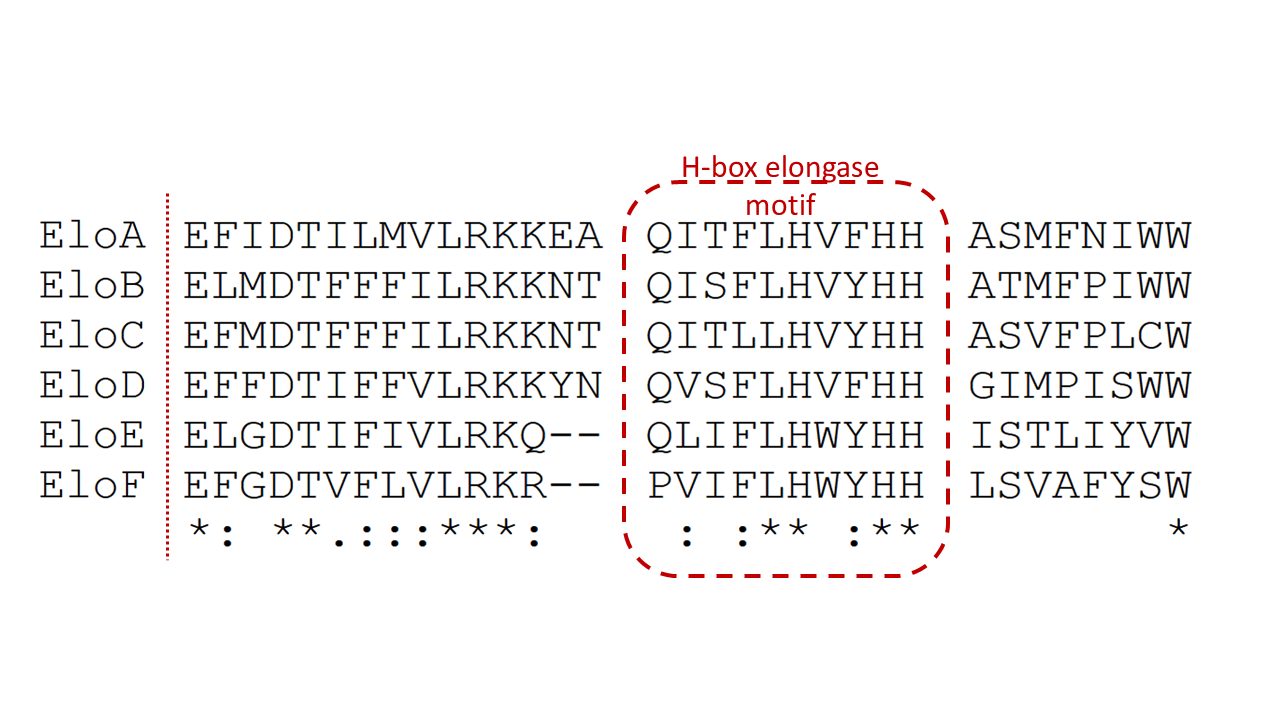

**Figure S1**. Alignment fragment of the deduced amino acid sequences of the *Platynereis dumerilii* elongase sequences assessed in this study (functional characterisation: EloA-D, only identified: EloE-F) aligned with Clustal Omega (default setting) as implemented in the software Geneious Prime (Sievers et al. 2011; Kearse et al. 2012). Elongase conserved motif containing the histidine box (HXXHH) and the upstream area to the -5 position (Hashimoto et al. 2008) is framed.

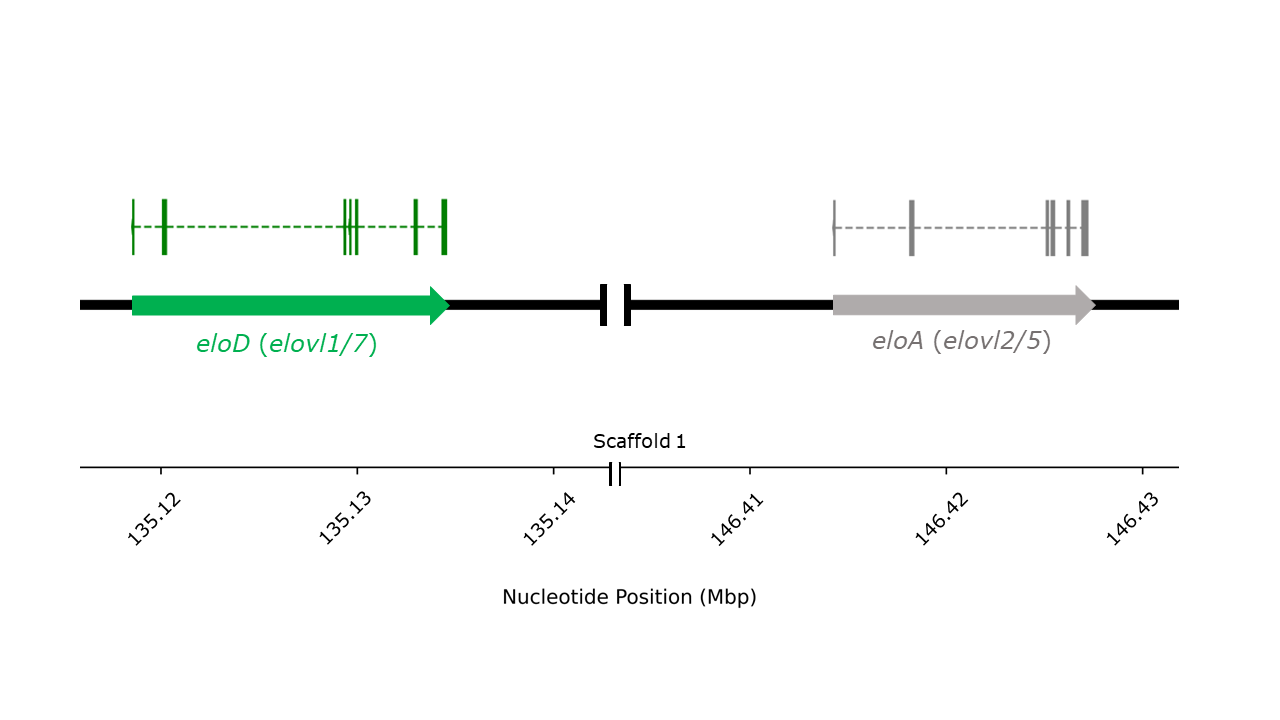

**Figure S2**. Gene structure comparison of *elovl1/7* and *elovl2/5* sequences on the same *Platynereis dumerilii* genomic locus (scaffold 1). This representation emphasises the conservation and variation in gene structure between the two homologues within the same scaffold.

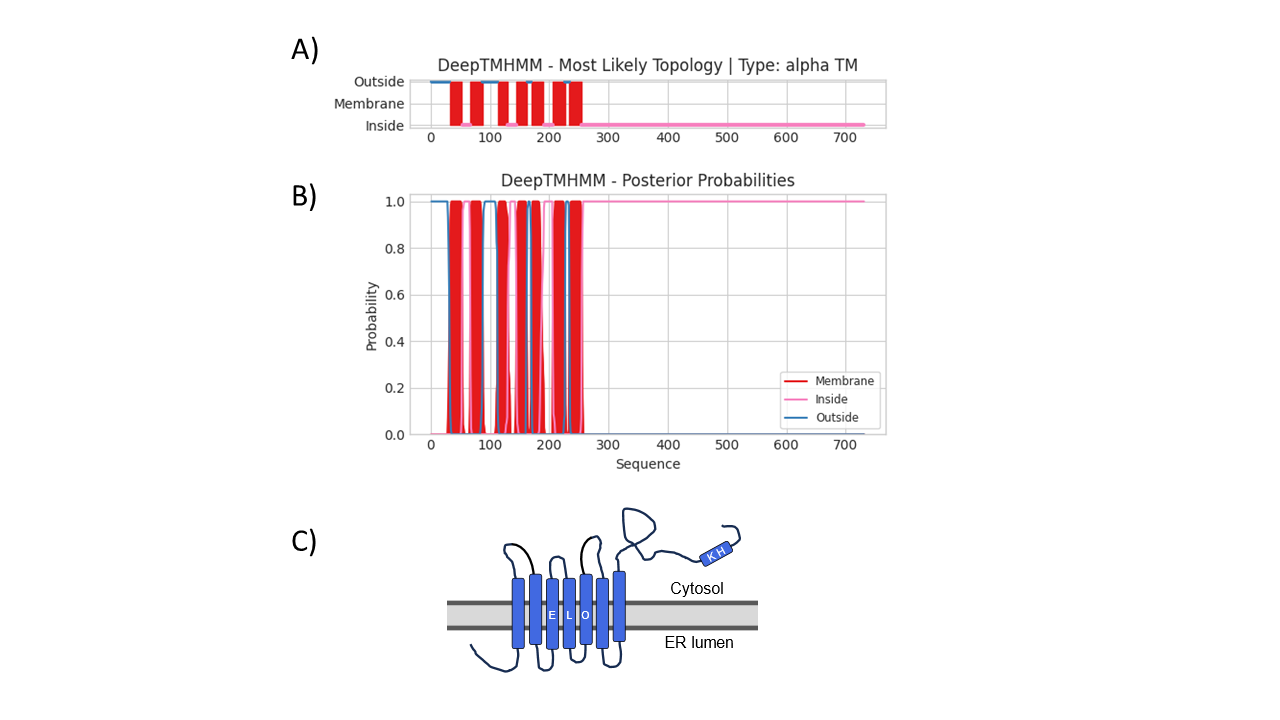

**Figure S3**. Deep learning prediction using DeepTMHMM (v.1.0.24; Hallgren et al. 2022) of the membrane topology of EloC (Elovl4): A) location of the amino acid chain with respect to its position in the membrane (inside/outside), or if it is a transmembrane section throughout the sequence (in amino acid position), and B) probability of predicted location. C) Scheme of the protein structure of EloC in the environment of the endoplasmic reticulum (ER). The functional predicted structures (Fig. 2A) are marked with coloured rectangles.

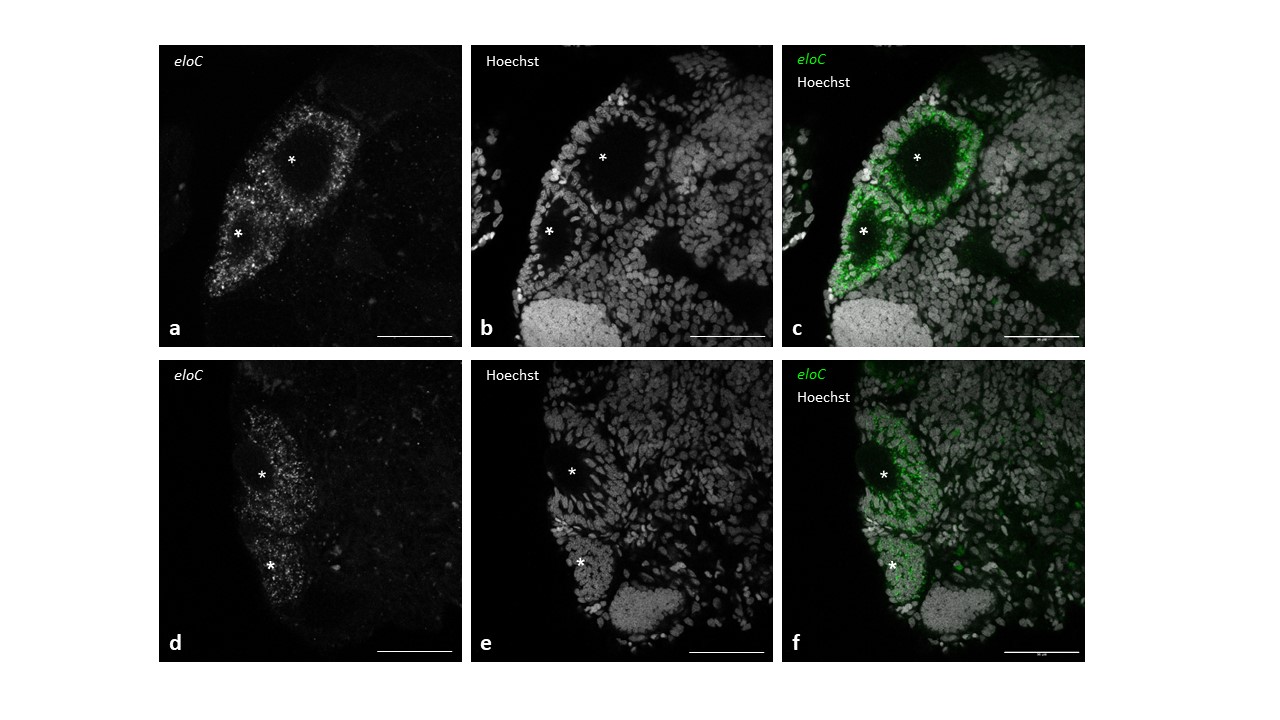

**Figure S4**. Visualisation, by in situ Hybridisation Chain Reaction (in situ HCR), of the expression of *eloC* in the *Platynereis dumerilii* head (**a, d**), Hoechst nucleic acid staining visualisation (**b, e**), and visualisation of *eloC* probes and Hoechst nucleic acid staining (**c, f**). Adult eyes are marked with asterisks. Acquisitions by confocal microscopy; scale bar = 50 μm.

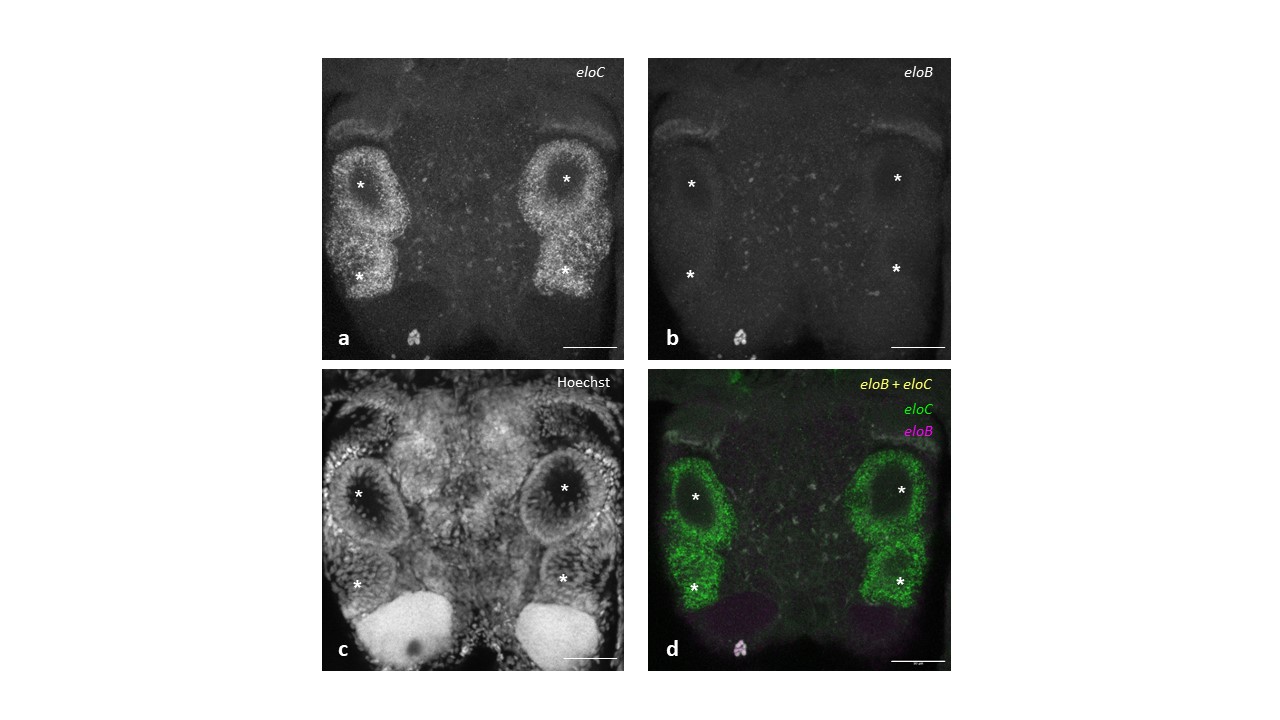

**Figure S5**. Visualisation of the expression of *eloB* and *eloC* in the *Platynereis dumerilii* head, by in situ Hybridisation Chain Reaction (in situ HCR): (**a**) Visualisation of *eloC* probes**,** (**b**) Visualisation of *eloB* probes, (**c**) Hoechst nucleic acid staining visualisation, and (**d**) Co-visualisation of *eloC* probes (green), *eloB* probes (magenta) and co-localisation of *eloB* and *eloC* probes (yellow). Adult eyes are marked with asterisks. Acquisitions by confocal microscopy; scale bar = 50 μm.

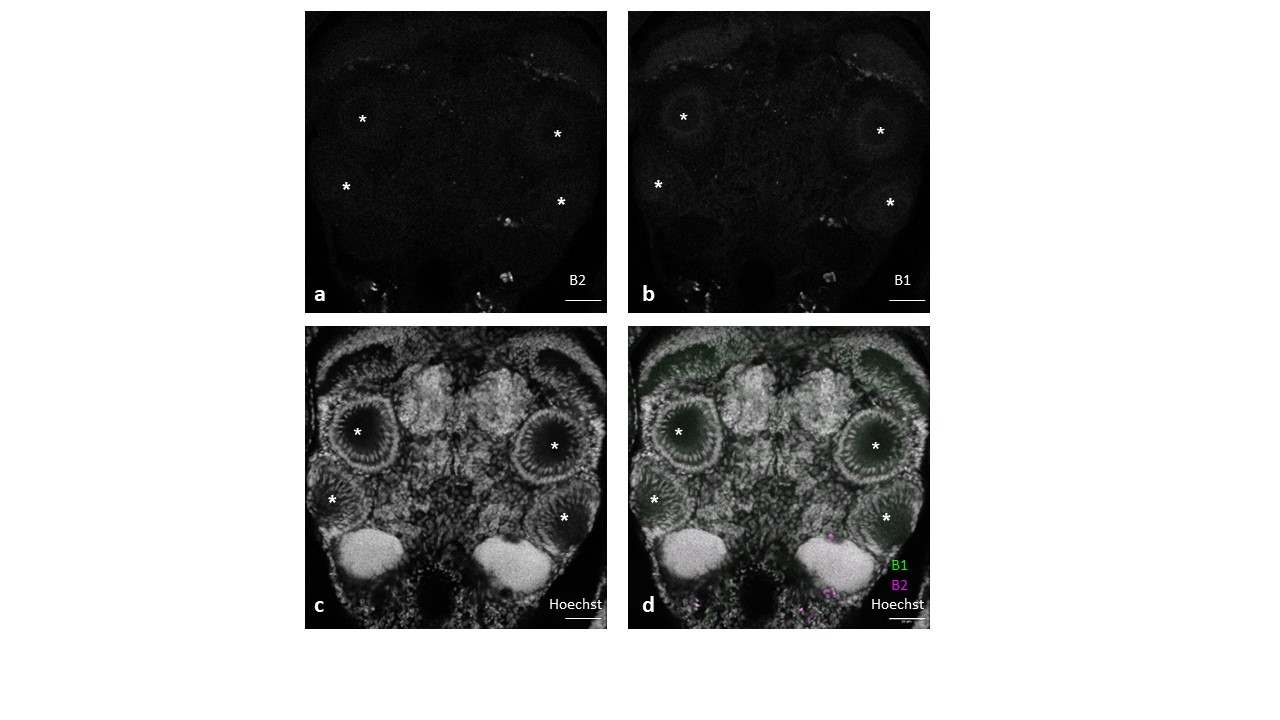

**Figure S6**. Control samples of *Platynereis dumerilii* heads were visualised using hybridisation chain reaction (HCR) for confocal microscope imaging. Samples were treated with HCR amplifier hairpins B1 and B2, respectively. (**a**) Control amplifier hairpins B2, (**b**) control amplifier hairpins B1, (**c**) Hoechst nucleic acid staining and (**d**) merged signals of amplifier hairpins B2 (magenta), amplifier hairpins B1 (green) and Hoechst nucleic acid staining (white). Adult eyes are marked with asterisks. Scale bar = 50 μm.

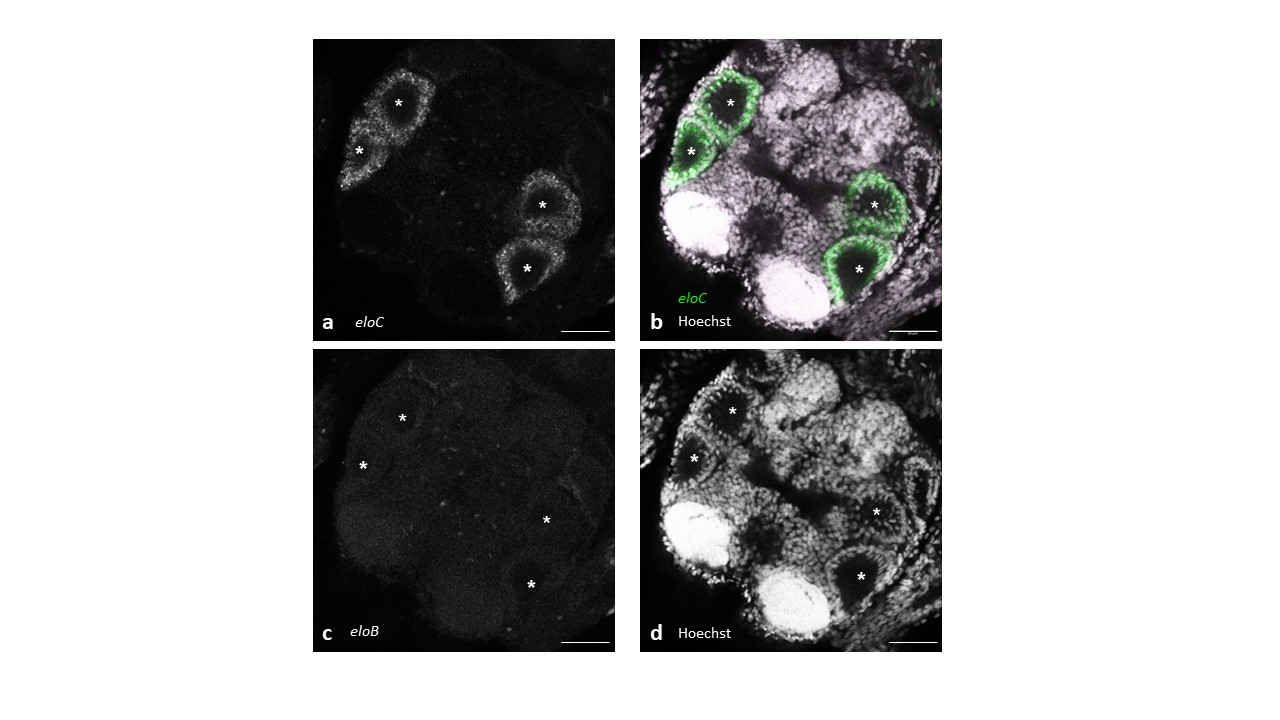

**Figure S7**. Visualisation of the expression of *eloC* in the *Platynereis dumerilii* head, by in situ Hybridisation Chain Reaction (in situ HCR): (**a**) Visualisation of *eloC* probes**,** (**b**) Visualisation of *eloC* probes plus Hoechst nucleic acid staining (white), (**c**) Visualisation of *eloB* probes, and (**d**) Hoechst nucleic acid staining (white). Adult eyes are marked with asterisks. Acquisitions by confocal microscopy; scale bar = 50 μm.
